# A billion-year-old bacterial machinery replicates plastid DNA and supports kleptoplastidy

**DOI:** 10.64898/2026.05.23.727376

**Authors:** Thibault Antoine, Fabien Burki, John M. Archibald, Eric Pelletier, Tom O. Delmont

**Author notes:** Corresponding authors: Correspondence should be addressed to Eric Pelletier and Tom. O. Delmont.

## Abstract

The endosymbiotic evolution of plastids and mitochondria was central to the origin and success of eukaryotes. One of the most prominent molecular machineries thought to have disappeared early in eukaryote evolution is the multi-subunit bacterial DNA polymerase III (DNApol-III), which is the principal enzyme complex supporting DNA replication in bacteria. Here, we combined worldwide metagenomics and cultivation to characterise the mosaic genomic landscape of abundant phytoplankton lineages of *Teleaulax* (Cryptophyceae), which contain an endosymbiotically-derived nucleomorph genome. Unexpectedly, the nuclear, plastid and nucleomorph genomes of *Teleaulax* contain ubiquitously expressed genes for plastid-targeted DNApol-III subunits. These genes shed light on the functioning of *Teleaulax* genomes when sequestered by the ciliate *Mesodinium* during its kleptoplastidic photosynthetic activity^1–3^. In particular, the alpha subunit gene (encoding the polymerase activity), which resides in the nucleomorph genome, is continuously expressed in *Mesodinium* in controlled laboratory experiments. This provides a mechanistic explanation for the replication of *Teleaulax* plastid genomes weeks after the nuclear genome is lost^4^. Beyond *Teleaulax* and close relatives, we also identified genes encoding plastid-targeted DNApol-III subunits (including alpha) in nuclear genomes of unicellular and multicellular lineages of Archaeplastida that form, along with those of Cryptophyceae, monophyletic clades firmly positioned within Cyanobacteria. Together, our results reveal a previously overlooked retention of bacterial DNA replication machinery from plastid primary endosymbiosis in Archaeplastida, its acquisition by Cryptophyceae during secondary endosymbiosis, and its direct role in contemporary plankton as a facilitator of kleptoplastidic photosynthetic activity by heterotrophic ciliates.

## Introduction

Photosynthetic eukaryotes have a considerable influence on global climate, planetary biogeochemical cycles, and the ecology of plankton in the sunlit oceans^5–9^. Their plastids all originated from a cyanobacterial endosymbiont in a heterotrophic ancestor of Archaeplastida, likely more than 1.5 billion years ago^10,11^. This rare event (the plastid primary endosymbiosis) represents the acquisition of photosynthesis by eukaryotes and directly resulted in the rise of Chloroplastida (green algae) as well as their land plant descendants, along with Rhodophyta (red algae) and Glaucophyta. After about 500 million years of photosynthesis in Archaeplastida, the plastids of green algae and red algae spread among distantly related eukaryotes by means of secondary and even tertiary endosymbiosis events^12^. In most cases, the endosymbiont’s nucleus was entirely lost. However, a few eukaryotic lineages have maintained an organelle containing a highly reduced remnant of this nucleus: the nucleomorph^13–15^. For instance, lineages within the Cryptophyceae (a class of algae prevalent in aquatic ecosystems) contain a nucleomorph of red algal origin^16–18^. Green and red algal-derived nucleomorphs have highly streamlined genomes*^17^* and shed light on the complex evolutionary trajectory of photosynthetic eukaryotes^18,19^.

Plastids contain a small genome compared to Bacteria, with many genes from the cyanobacterial endosymbiont having been transferred to the nuclear genome. These genes encode proteins that are translocated into the plastid organelle and contribute to its functioning^12,20–23^. In the case of secondary and tertiary endosymbiosis events, the nucleomorph genome also produces plastid-targeted proteins of cyanobacterial origin that act in concert with those of the nuclear and plastid genomes. Yet, many cyanobacterial core genes have never been observed in eukaryotes, indicating that they were lost during the plastid primary endosymbiotic event. A prominent molecular machinery thought to have been lost during the symbiont-to-plastid transition is that of DNA polymerase III (DNApol-III), which includes multiple subunits and is the principal enzyme complex involved in bacterial DNA replication^24,25^. In photosynthetic eukaryotes, replication of the plastid genome is performed by nucleus-encoded polymerases labelled as “plant organellar DNA polymerases” (POP)^26–28^. POP has an evolutionarily origin unrelated to that of DNApol-III^29,30^.

The last decade has seen a rapid increase in environmental genomic databases for the study of marine plankton, which includes nuclear and plastid genomes from unicellular photosynthetic eukaryotes prevalent in the sunlit oceans^9,31–33^. Here, we took advantage of *Tara* Oceans metagenomes^34^ to perform the first genome-resolved metagenomic survey of nucleomorphs. Our data contain several high-quality nucleomorph genomes from the abundant and common Cryptophyceae genus *Teleaulax*. We subsequently resolved the triumvirate environmental genomes (nuclear, nucleomorph, plastid) of multiple *Teleaulax* lineages. Prior to our survey, only one *Teleaulax* plastid had been characterised^35^. Unexpectedly, we found that *Teleaulax* lineages (including a cultured species) contain and express central DNApol-III subunits for bacterial DNA replication, spread across the nuclear, nucleomorph and plastid genomes. The most important subunit alpha (ensuring polymerase activity) occurs in the nucleomorph and is predicted to translocate into the plastid organelle. We investigated the ecological prominence of these DNApol-III subunits in the context of kleptoplastidic photosynthetic activity^4,36,37^ and traced their evolutionary journey all the way back to plastid primary endosymbiosis.

## Results

### A genome-resolved metagenomic survey of marine nucleomorphs

As part of global efforts characterising plankton genomics in the sunlit oceans, large metagenomic co-assemblies have previously been obtained from 280 billion metagenomic reads of the *Tara* Oceans expeditions^31,34^ (Table S1). These assemblies contain thousands of metagenome-assembled genomes (MAGs) of Bacteria, Archaea, as well as the nuclear and plastid genomes of eukaryotes, and associated DNA viruses^9,31,38,39^. Here, we used a subunit of the DNA–dependent RNA polymerase as a focal marker gene to screen for planktonic nucleomorphs (see Methods). We identified a eukaryotic clade corresponding to Cryptophyceae nucleomorphs and used this phylogenetic signal as guidance (as in refs^9,39^) to perform the first genome-resolved metagenomic survey of nucleomorphs. Using the anvi’o bioinformatic platform^40,41^, we characterised and manually curated nine MAGs of nucleomorphs from co-assemblies of the Atlantic Ocean (n=4), Pacific Ocean (n=3) and Arctic Ocean (n=2). These nucleomorph MAGs have an average size of 466 kb (up to 602 kb) and an average completion of 79% (Table S2). They correspond to seven distinct Cryptophyceae lineages (average nucleotide identity (ANI) below 98%) that share a relatively large core functional pool covering various ribosomal and plastid-associated proteins (Table S2).

No nucleomorph genomes are available from environmental uncultured phytoplankton^16–18,42–44^ (Table S2). For Cryptophyceae cultured strains, nucleomorph genomes are available for the Cryptomonadales (n=8) and Pyrenomonadales (n=1; *Guillardia theta*). These genomes average ∼590 kb in length and share many genes with those of our nucleomorph MAGs (Table S2). We used a concatenation of 33 single copy core genes (see Methods) to perform a multi-gene phylogenetic analysis including the 9 nucleomorph MAGs, the 9 available red algal-derived nucleomorphs from cultured strains as well as representatives of red algae (Table S2). The nucleomorph MAGs form a well-supported clade sister to *Guillardia theta* within the order Pyrenomonadales (Figure 1). Among the *Tara* Oceans metagenomes, nucleomorph MAGs mainly occur in the 0.2-3/0.8-5 µm size fraction and are enriched in the surface layer compared to the deep chlorophyll maximal one (Table S3 and Figure S1). Their transcripts are also abundant in the corresponding metatranscriptomes (Table S4). In contrast, nucleomorph genomes similar to those of cultured strains were undetected in these *Tara* Oceans samples (Table S3). Together, the nucleomorph MAGs correspond to relatively small (below 5 µm) but abundant and transcriptionally active eukaryotic cells (Figure 1, Table S3), which based on our environmental genomic linkage (see next section), are affiliated to the genus *Teleaulax*.

**Figure 1:**
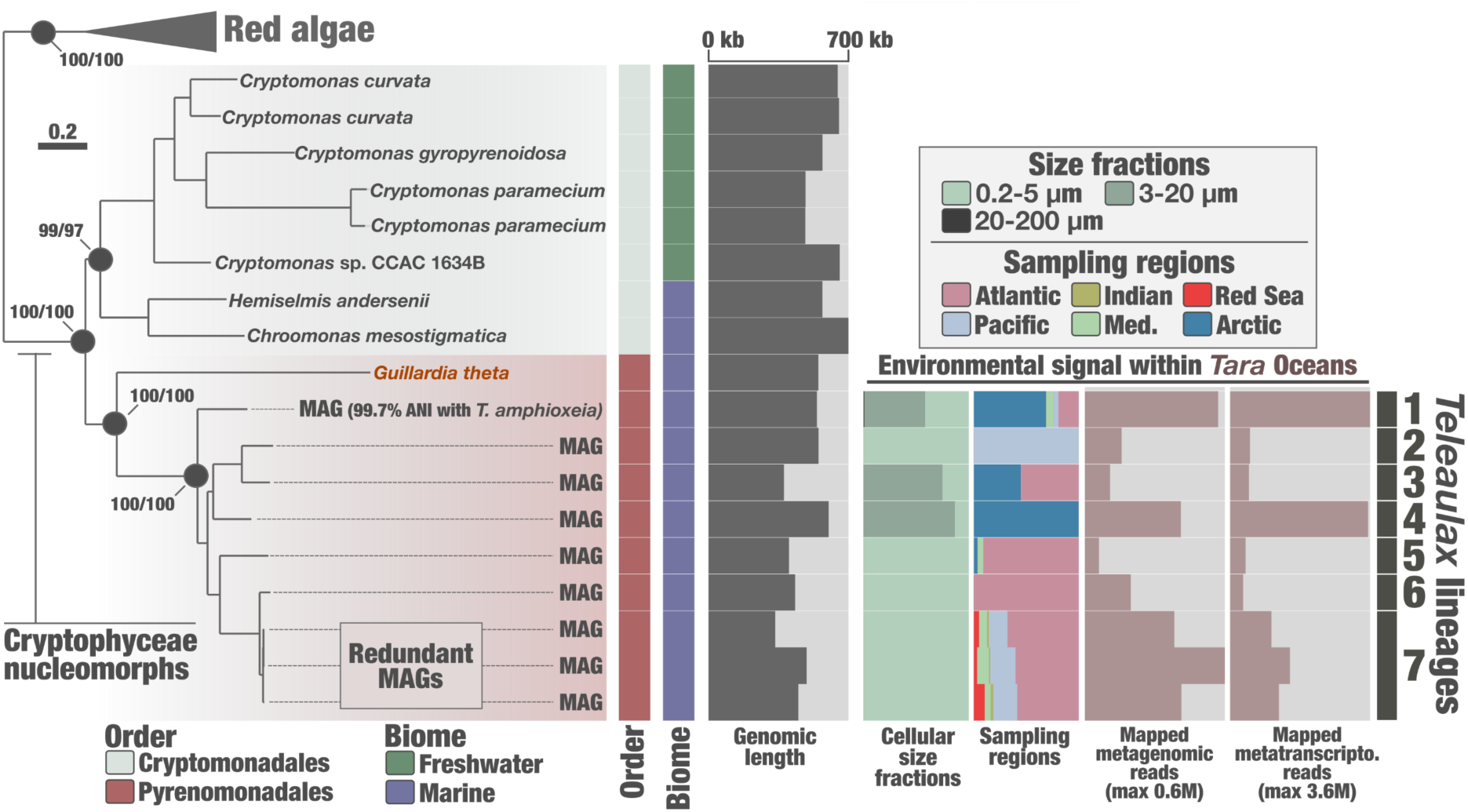
Phylogeny of red algal-derived nucleomorphs. At left is a maximum-likelihood phylogenetic tree encompassing eight red algal nuclear genomes as well as 18 cryptophyte nucleomorph genomes derived from metagenomics and cultured strains. The tree is based on a concatenation of 33 single-copy core genes (6,763 amino acid positions) and was built using IQ-TREE with the LG+F+R5 model. The tree is rooted between red algal nuclear and nucleomorph genomes. Black dots indicate nodes with high support, corresponding to Shimodaira–Hasegawa-like approximate likelihood ratio test (SH-like aLRT, left score) and ultrafast bootstrap (UFBoot; right score). The tree shows additional annotation layers. The cellular size fraction and sampling region signal is based on metagenomics. The 0.2-5 µm size fraction encapsulates the 0.2-3 µm and 0.8-5 µm size fractions, while the 3-20 µm size fraction encapsulates the 3-20 µm and 5-20 µm size fractions of *Tara* Oceans samples. The scale bar represents 0.2 substitutions per site.

### Resolving the environmental genomes of prevalent *Teleaulax* lineages

To gain further knowledge on the nucleomorph MAGs, we characterised the nuclear and plastid genomes also present in the same cells. To do so, we incorporated nucleomorph MAGs into a planktonic eukaryotic genomic database that includes 683 marine nuclear MAGs^31^, 705 marine plastid MAGs and 163 plastid genomes from cultured strains^9^ (Table S5). In this database, eight nuclear genomes and 25 plastid genomes correspond to Cryptophyceae. Our main rationale was that the relative abundance of species-level genomes (nuclear, nucleomorph, plastid) should strongly correlate across *Tara* Oceans samples, and that phylogenetic inferences would support this environmental genomic linkage. For each nucleomorph MAG, we performed a Pearson correlation of coverage distribution across 1,178 *Tara* Oceans metagenomes against those of each genome of the planktonic eukaryotic genomic database (see Methods).

As expected, most genomes in the database poorly correlate with the nucleomorph MAGs (average Pearson correlation of 0.03 when considering all pairs). However, the nucleomorph MAGs strongly correlate with Cryptophyceae nuclear MAGs (n=7), Cryptophyceae plastid MAGs (n=5), and one plastid genome^35^ of the cultured *Teleaulax amphioxeia* strain HACCP-CR01, with an average Pearson correlation of 0.91 across these 13 pairs (Figure 2 and Table S3). With this approach, four out of the seven non-redundant nucleomorph MAGs were unambiguously linked to both a nuclear and a plastid MAG. An additional nucleomorph MAG was linked to a nuclear MAG and the *Teleaulax amphioxeia* plastid genome^35^. The abundance correlation was further supported by the congruent phylogenies recovered from the three types of genomes, where MAGs all grouped with *Guillardia theta^9,31^* (Figure 2). Metagenomic signals indicate that, for each lineage, the nuclear and nucleomorph genomic copy numbers were almost identical in the environment while the plastid genome displayed much higher copy numbers (from ∼5X to ∼80X more coverage per sample depending on the lineage) (Table S3). Furthermore, the plastid genome sequenced from the cultured *Teleaulax* allowed to confidently assign the nucleomorph MAGs to this genus of Pyrenomonadales. *Teleaulax* emerges as one of the most prevalent nucleomorph-containing lineages in the upper oceans.

**Figure 2:**
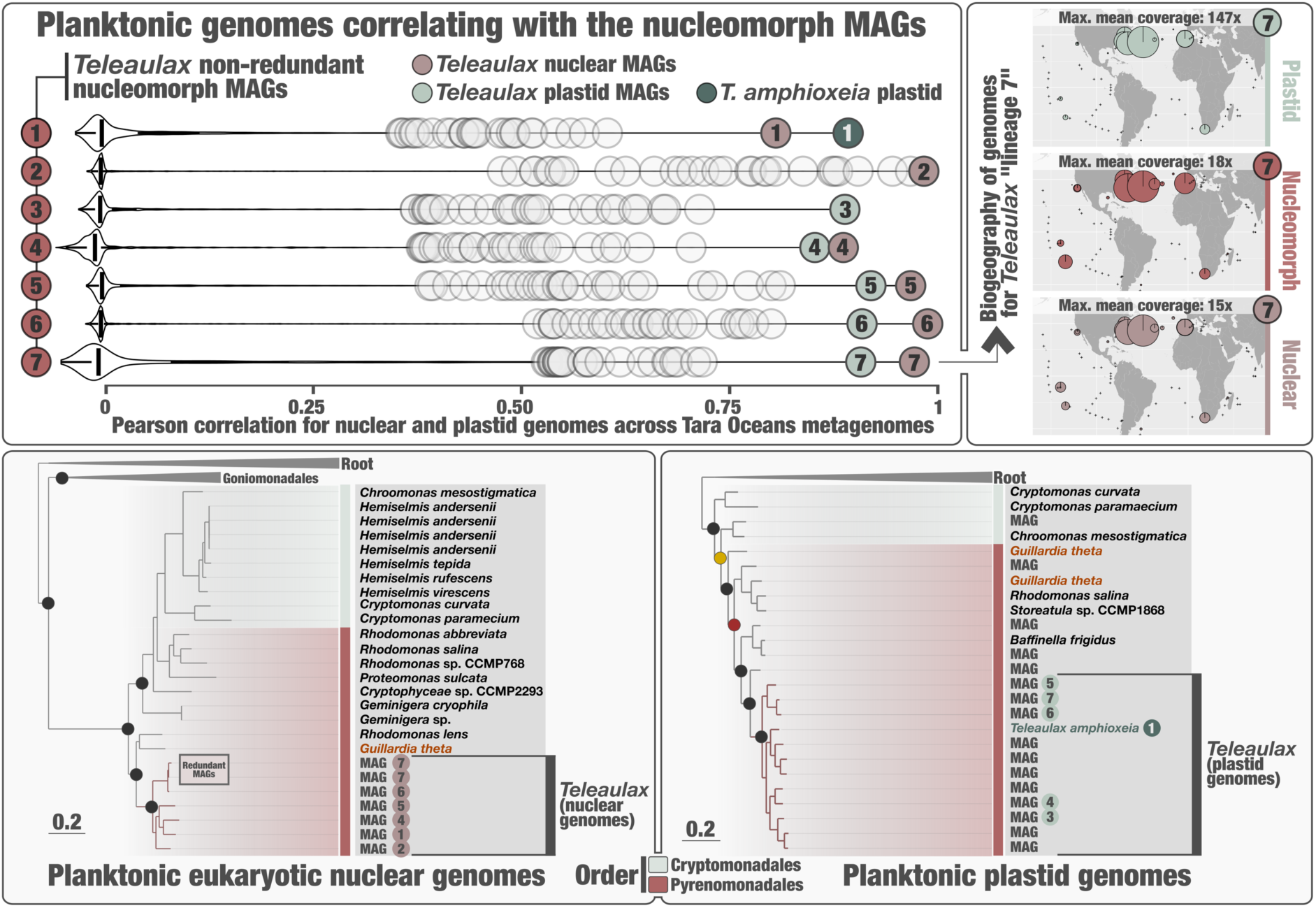
Environmental genomic linkage of *Teleaulax* lineages. The top left panel displays, as a violin plot for each of the nucleomorph non-redundant MAGs (n=7), the Pearson correlation coefficients distributions of MAG mean coverage with that of each genome of the planktonic eukaryotic genomic database (n=1,553) across 1,178 *Tara* Oceans metagenomes. Nuclear and plastid genomes strongly correlating with nucleomorph MAGs and affiliated to Cryptophyceae are highlighted in red and green, respectively. Only the 30 highest Pearson correlation coefficients are visualised as a circle for each violin plot. The top right panel displays the relative abundance of nuclear, nucleomorph and plastid genomes of one *Teleaulax* lineage in the sunlit ocean, with the size of dots in the word maps corresponding with the mean genomic coverage across *Tara* Oceans metagenomes of the 0.2-5 µm size fraction. Bottom panels display global phylogenies^9,31^ for the nuclear (left panel; 7,243 amino acid positions; LG+F+R10 model; 576 sequences) and plastid genomes (right panel; LG+F+I+G4 model; 839 sequences) of the planktonic eukaryotic genomic database (all lineages outside of Cryptophyceae are used for rooting), which highlight the *Teleaulax* genomes linked based on Pearson correlation coefficients to a nucleomorph MAG (same numbering as in top panels). In the bottom left panel, branch support highlighted dots (see Methods for more details) were considered high (SH-like aLRT ≥ 80 and UFBoot≥ 95, in black). In the right panel, supports were considered high (UFBoot≥ 95, in black), medium (UFBoot 80-95, in yellow) or low (UFBoot<80, in red). Scale bars indicate the number of substitutions per site for each tree.

We sequenced the genomes of *T. amphioxeia* strain AND-A0710 to further ground insights derived from our metagenomic investigations. A combination of PacBio and Hi-C sequencing provided a telomere-to-telomere assembly of the nuclear genome (262 Mb organised in 101 chromosome-scale pseudo-chromosomes), nucleomorph genome (560 kb) and plastid genome (130 kb) (Figure S2 and Table S6). *T. amphioxeia* strain AND-A0710 corresponds to one of the seven lineages characterised from marine metagenomes, with a near-perfect match for the nuclear genome (99.43% ANI, the MAG being 26.74% complete) and nucleomorph genome (99.67% ANI, the MAG being 85.75% complete) (Table S2), supporting the quality of Cryptophyceae nuclear and nucleomorph MAGs.

### A bacterial machinery for plastid DNA replication spread across three genomes in *Teleaulax*

Our extended database of Cryptophyceae nucleomorph genomes contains a large functional core shared among lineages of the orders Cryptomonadales and Pyrenomonadales, including several genes encoding plastid-associated proteins^18,19^ (Table S2). More surprisingly, *Teleaulax* nucleomorph genomes also contain a gene of nearly 4 kb encoding DNApol-III alpha subunit, which is missing in the other cryptophyte clades and has not been documented in eukaryotes except in *Paulinella*^45,46^. Expanding our investigation to other *Teleaulax* genomes, we identified additional DNApol-III subunit genes for epsilon (exonuclease activity) and beta (sliding clamp) in the nuclear genomes and tau/gamma (clamp loader complex) in the plastid genomes (Table S5). Whereas the tau/gamma subunit was previously found in *Pyrenomonadales* plastid genomes*^47^*, this is the first report of alpha and beta subunits in eukaryotes. Genomic data from *Teleaulax amphioxeia* validate these results (Table S5) while metatranscriptomics from *Tara* Oceans indicates that the DNApol-III subunits are expressed by all seven *Teleaulax* lineages (Table S4).

Given its central role in bacterial DNA replication, we hypothesized that DNApol-III, and especially the polymerase activity of the alpha subunit, might support plastid genome replication in *Teleaulax*. To determine whether all subunits can act in concert in the plastid, we searched for targeting signals in their amino acid sequences that could specify organelle targeting (see Methods). We found amino (N)-terminal plastid transit peptides in the alpha and beta subunits (Figure S3 and Table S7). In particular, the beta subunit contains a clear bipartite plastid targeting signal consistent with a transit from the cytoplasm to a secondarily-derived, multi-membrane bound plastid via the endomembrane system^48^. A potential plastid transit peptide (weaker signal compared to alpha and beta) was also found on the N-terminus of the epsilon subunit. Overall, the occurrence of tau/gamma in the *Teleaulax* plastid genome and putative plastid transit peptides on alpha (nucleomorph genome) and beta (nuclear genome) subunits strongly suggest that DNApol-III includes plastid-associated proteins and replicates plastid DNA in *Teleaulax*.

### Plastid genomic replication during *Teleaulax*-*Mesodinium* kleptoplastidic activity

*Teleaulax* is not only a cosmopolitan and abundant member of marine phytoplankton communities, but also the preferred prey and plastid source of the kleptoplastidic ciliate *Mesodinium^37^.* This plastid symbiosis can fix carbon for up to two months, spanning multiple cell division cycles owing to a tightly regulated plastid sequestration^36,49^. Inside the ciliate, *Teleaulax*-derived, membrane-delineated plastid-nucleomorph-mitochondrion complexes (hereafter named “sequestered organelle complexes”^50^) self-replicate and support photosynthesis activity even after the *Teleaulax* nuclear genome (and the POP polymerases it encodes) is lost^36,51,52^. To follow the expression of *Teleaulax* DNApol-III subunits inside *Mesodinium*, and to screen for the occurrence of complementary *Mesodinium*-encoded DNApol-III subunits, we combined our new genomic data of *T. amphioxeia* strain AND-A0710 with an available RNA sequencing data set covering the kleptoplastidic photosynthetic activity of *Mesodinium rubrum* after ingestion of *T. amphioxeia* in controlled laboratory conditions^51^.

First, we identified a DNApol-III epsilon subunit among the *Mesodinium*-derived transcripts. This gene is evolutionary unrelated to that of *Teleaulax* (Figure S4), is expressed across multiple time points (Table S8), and shows a significant increase in transcriptomic signal in the late phase (induced starvation) compared to the early phase (when *Mesodinium* is well fed with free-living *Teleaulax* cells) of the kleptoplastidic activity (t-test; p-value of 10^-6^). We did not find any clear signal for a plastid transit peptide on this transcript. In addition, no other DNApol-III subunits were identified among the *Mesodinium* transcripts. Second, we found that the gene encoding the alpha subunit of *Teleaulax* is continuously expressed throughout the experiment, including during the late phase of the kleptoplastidic activity (Table S8 and Figure S5). The genes encoding the beta and epsilon subunits of *Teleaulax* are also continuously expressed but show a clear spike (∼17x enrichment) for the beta subunit after ingestion by *Mesodinium* compared to the free-living form (t-test; p-value of 1.7.10^-22^). These results suggest that DNApol-III is involved in plastid DNA replication not just in free-living *Teleaulax* cells but also inside the sequestered organelle complex in *Mesodinium*, a process that is possibly augmented by a ciliate-derived subunit (epsilon).

### The bacterial DNA replication machinery originates from plastid primary endosymbiosis

To assess whether DNApol-III subunits are present in other eukaryotes beside *Teleaulax*, we searched for homologs in the planktonic eukaryotic genomic dataset (environmental genomics) and eukaryotic genomic data from NCBI (mostly cultured strains) (see Methods) (Figure 3). We identified genes encoding the DNApol-III epsilon subunit in a wide range of eukaryotic lineages (Tables S5 and S9). In contrast, we only identified genes encoding the alpha and tau/gamma subunits in Pyrenomonadales genomes (with alpha and tau/gamma genes always occurring in the nucleomorph genome and plastid genome, respectively) and in nuclear genomes of unicellular and multicellular lineages of Archaeplastida (Table S5 and Table S9). We found that the alpha subunit occurs more commonly in red algae compared to green algae, where it is systematically missing in Chlorophyta (Figure S6). Finally, the beta subunit appears limited to the Pyrenomonadales. Congruent with *Teleaulax*, we found clear signal for N-terminal plastid transit peptides on the alpha and beta subunits of the other Pyrenomonadales (Table S7), as well as on the alpha and tau/gamma encoded in the nuclear genomes of Archaeplastida (Figure S7). Those results indicate that DNApol-III occurs in plastid-targeted proteins in two distantly related eukaryotic lineages, Pyrenomonadales and Archaeplastida, but which are linked by endosymbiotic history.

**Figure 3:**
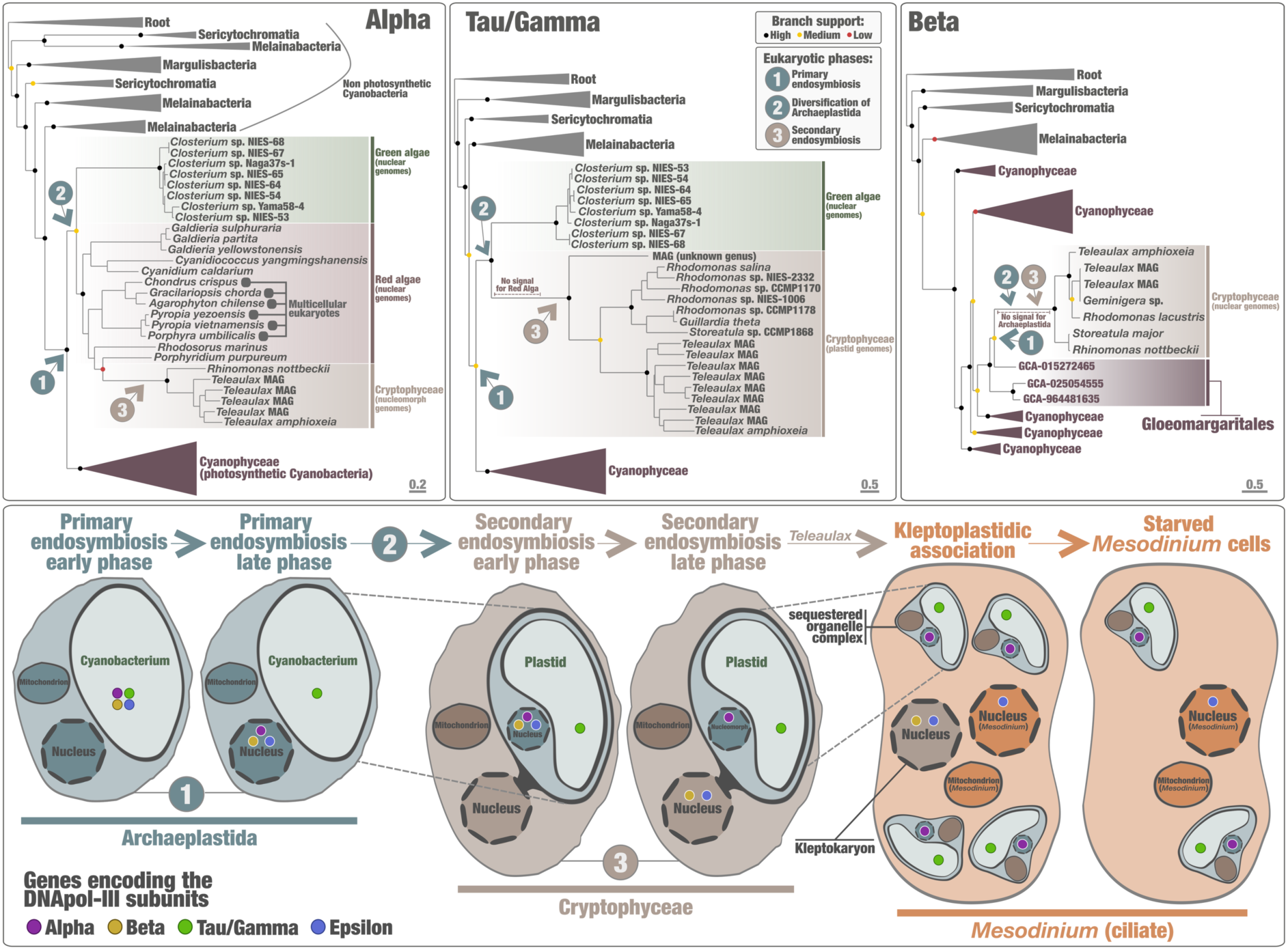
Evolutionary trajectory of cyanobacterial-derived DNA polymerase machinery among photosynthetic eukaryotes. Left panel displays a maximum-likelihood phylogenetic tree of DNApol-III alpha subunit (1,179 amino acid positions; LG+R10 model) that includes 567 sequences (495 Cyanobacteria, 27 Eukarya, 17 Chloroflexia and 28 Actinomycetota). Middle panel displays a maximum-likelihood phylogenetic tree of DNApol-III tau/gamma subunit (695 amino acid positions; Q.PFAM+F+R10 model) that includes 450 sequences (397 Cyanobacteria, 25 Eukarya, and 28 Actinomycetota). Right panel displays a maximum-likelihood phylogenetic tree of DNApol-III beta subunit (391 amino acid positions; LG+F+R9 model) that includes 505 sequences (454 Cyanobacteria, 7 Eukarya, 16 Chloroflexia and 28 Actinomycetota). Alpha and beta trees were rooted with Chloroflexia and Actinomycetota, and the tau/gamma tree was rooted with Actinomycetota. In addition, highlighted phylogenetic supports (dots in the trees) were considered high (SH-like aLRT ≥ 80 and UFBoot ≥ 95, in black), medium (SH-like aLRT ≥ 80 or UFBoot ≥ 95, in yellow) or low (SH-like aLRT < 80 and UFBoot < 95, in red). Scale bars indicate the number of substitutions per site for each tree. Trees were decorated with additional information and visualised with anvi’o. Bottom panel summarises an evolutionary model for the DNApol-III subunits from primary endosymbiosis to contemporary kleptoplastidic photosynthetic activity within the ciliate *Mesodinium*.

The phylogenetic relationships of these subunits among eukaryotes recapitulate the expected branching patterns resulting from successive endosymbiosis (Figure 3). Cryptophytes are most closely related to red algae in the alpha subunit phylogeny, and sister the green algae in the tau/gamma subunit (red algae missing) phylogeny. In all cases, the cryptophyte-archaeplastid monophyletic group (or just cryptophytes for the beta subunit) are firmly positioned inside Cyanobacteria, as a sister clade to the class Cyanophyceae that encapsulates all photosynthetic cyanobacteria for alpha and tau/gamma, or nested within this class and most closely related to Gloeomargaritales for beta (Figure 3, Figure S8 and Table S10). Taken together, these phylogenies indicate that the DNApol-III subunits were introduced in Archaeplastida from Cyanobacteria during primary endosymbiosis and later transferred to cryptophytes from red algae during secondary endosymbiosis. Our results expose a remarkably long evolutionary journey of a cyanobacterial DNA replication machinery from plastid primary endosymbiosis to contemporary photosynthetic eukaryotes, with *Mesodinium* relying on this relic to sustain its kleptoplastidic activity after ingestion of *Teleaulax* cells.

## Discussion

*Teleaulax* occurs broadly in the oceans, where it is involved in the carbon cycle both as free-living cells and as plastid donor in the kleptoplastidic relationship with the ciliate *Mesodinium*^1–3^. Like most Cryptophyceae, *Teleaulax* contains a red algal-derived nucleomorph^14–16,42^, which supports plastid functioning. Prior to our survey, the only documented genomic characterisation of *Teleaulax* was that of a plastid genome from culture^35^. Using genome-resolved metagenomics (*Tara* Ocean metagenomic data^34,53^) and phylogeny-aware environmental genomic linkage, we characterised environmental nucleomorph genomes and subsequently resolved the nuclear, nucleomorph and plastid genomes of prevalent *Teleaulax* lineages, which include sequences closely related to the cultured model species *Teleaulax amphioxeia*. Our complete genomic characterisation of *Teleaulax amphioxeia* strain AND-A0710 validated this environmental genomic survey and supported our investigations of the ecological role and evolutionary prominence of DNApol-III subunits in *Teleaulax* and other photosynthetic eukaryotes.

Present-day expression of DNApol-III subunits by all characterised *Teleaulax* lineages, as inferred from global scale metatranscriptomics, attests to the active utilisation of a bacterial DNA replication machinery relic in prevalent marine photosynthetic unicellular eukaryotes. With the tau/gamma subunit encoded in the plastid genome, and alpha (nucleomorph genome) and beta (nuclear genome) subunits predicted to translocate into the plastid organelle, these DNApol-III subunits appear to directly support plastid genomic replication. To our knowledge, this is the first documentation of gene complementarity across three genomes (nuclear, nucleomorph and plastid) of the same species to perform a fundamental cellular process. Beyond the role of DNApol-III subunits in free-living *Teleaulax* cells, this ‘three-genomes’ functional architecture sheds an unexpected light into *Mesodinium* kleptoplastidic activity. Indeed, *Teleaulax* plastids are preferentially sequestered alongside the nucleomorph and mitochondria (the sequestered organelle complex) inside *Mesodinium* cells and support multi-generational photosynthetic activity^37,51^. The *Teleaulax* nucleomorph was previously hypothesised to contribute to plastid genomic replication using an unspecified mechanism^51^. We have shown that the nucleomorph-encoded DNApol-III alpha subunit encodes an N-terminal plastid transit peptide predicted to target to the plastid; the gene is expressed in free living *Teleaulax* cells and when they are sequestered by *Mesodinium*. This might explain how plastid genome replication persists after the *Teleaulax* nuclear genome is lost during the late phase of this kleptoplastidy. Together with laboratory-based experiments (e.g., ^36,49^), the genomic data presented here for *Teleaulax* support the hypothesis that DNApol-III subunits contribute to kleptoplastidic stability in *Mesodinium* by facilitating plastid genome replication.

Except in the case of a second primary endosymbiosis event (∼100 million year ago^54^) in the amoeboid protist *Paulinella*^45,46^, the central component of the cyanobacterial DNA replication machinery (i.e., DNApol-III alpha subunit) has not been documented in eukaryotes. Other cyanobacterial DNApol-III subunits were also considered to have been lost during or soon after the origin of canonical primary plastids more than one billion years ago, owing to their absence in previous genomic data of photosynthetic eukaryotes. In this context, our identification of cyanobacterial DNApol-III subunits scattered in the nuclear (beta, epsilon), nucleomorph (alpha) and plastid (tau/gamma) environmental genomes of *Teleaulax*, with strain AND-A0710 confirming their occurrence and spread across the genomes, was unexpected. The occurrence of these subunits in additional Pyrenomonadales genera as well as in a subset of unicellular and multicellular lineages of Archaeplastida (both green algae and red algae) exposed a long evolutionary history among eukaryotes. Indeed, eukaryotic genes encoding the alpha, beta and tau/gamma subunits form monophyletic groups firmly positioned within Cyanobacteria. The evolutionary relationship of the alpha subunit in eukaryotes reflects the expected history from primary and secondary endosymbiosis events. In addition, the presence of beta subunit (constrained to Pyrenomonadales within the Cryptophyceae) links *Teleaulax* and closely related genera to *Gloeomargaritales* well inside the photosynthetic cyanobacteria. With *Gloeomargaritales* previously suggested to be the closest known lineage to plastids^55^, the evolution of DNApol-III beta is consistent with this evolutionary relationship. Overall, genes encoding these DNApol-III subunits in eukaryotes support the previously overlooked maintenance of a cyanobacterial DNA replication machinery from primary endosymbiosis in a wide range of Archaeplastida and its acquisition by the common ancestor of Cryptophyceae from a red alga via secondary endosymbiosis.

Together, the occurrence of DNApol-III subunits among eukaryotes and their phylogenetic signal is best explained by a model spanning more than one billion years of evolution (Figure 3). First, DNApol-III subunits were maintained in the Archaeplastida host lineage involved in the endosymbiont-to-plastid transition phase of primary endosymbiosis to support genomic replication of the emerging plastid (phase I). DNApol-III subunits (some having been transferred to the host nuclear genome) remained in lineages of green algae and red algae following the Archaeplastida diversification into three major high-ranking taxonomic lineages^54,56^ (phase II). This phase spans the period of photosynthetic activity constrained to Archaeplastida among eukaryotes. We did not identify any present-day lineages of Archaeplastida containing all four DNApol-III subunits, with only a few sporadic remnants attesting to phase II. Subsequently, red algal DNApol-III subunits were acquired by the ancestors of Cryptophyceae during a second endosymbiosis event, and some subunits were transferred from the red algal nuclear genome to the Cryptophyceae nuclear genome, this time during or after the reductive evolution that resulted in nucleomorphs (phase III). Present-day *Teleaulax* lineages are widespread and abundant marine representatives of this phase, attesting to the long-lasting evolutionary journey of DNApol-III subunits among the eukaryotes. Finally, several Cryptophyceae lineages (and most notably the order Cryptomonadales) lost those DNApol-III subunits (phase IV). It is unclear when the vast majority of Archaeplastida lineages fully or partially lost the DNApol-III subunits, and if other photosynthetic eukaryotic clades (e.g., the stramenopiles and haptista) ever acquired the bacterial DNA replication machinery by means of secondary and higher-level endosymbiosis events, nor why *Teleaulax* and its close relatives within Pyrenomonadales are thus far the only known eukaryotes (apart from *Paulinella*) that maintain DNApol-III beta subunit. The fact that *Mesodinium* preferentially preys on Pyrenomonadales but not on its sister order (Cryptomonadales) is particularly revealing. Far from a complete loss following plastid acquisition, we conclude that DNApol-III has a long record of vertical evolution among photosynthetic eukaryotes (with various gene loss events) and plays an active role in ecologically prominent kleptoplastidic photosynthetic activities within plankton.

## Methods

### *Tara* Oceans metagenomes and metatranscriptomes

We analyzed 1,178 metagenomes and 1,165 metatranscriptomes from the *Tara* Oceans available under project accession PRJEB402 at the EBI. Summary on the metagenomes and metatranscriptomes can be found in Table S1 (including the number of reads and environmental metadata).

### Phylogeny-guided genome-resolved metagenomics for nucleomorphs

We used two complementary datasets (metagenomic assemblies and evolutionary-informative proteins) to characterise nucleomorph MAGs. On the one side, we used 11 large metagenomic co-assemblies from *Tara* Oceans (∼12 million contigs) which were organised into 2,550 metabins using constrained automatic binning and processed with anvi’o^40,41^ v.8 for manual binning purposes. On the other side, we used proteins corresponding to the DNA-dependent RNA polymerase A subunit (RNApolA) and found in those *Tara* Oceans metabins (those were identified using HMMER^57^ v3.1b2), in the context of a phylogeny of representative sequences (amino acid level with sequence similarity <90%). We were intrigued by a small RNApolA clade basal to that of all eukaryotes, for which NCBI blast results pointed towards previously uncharacterised nucleomorphs of red algae origin. We performed a targeted genome-resolved metagenomics (using the RNApolA clade as guidance) to characterise the corresponding environmental genomes, as previously done among the same data sets for large and giant eukaryotic DNA viruses^39,58^ as well as plastids^9^. Briefly, we manually inspected metabins containing the focal RNApolA clade using the anvi’o interactive interface, in which contigs are organised based on sequence composition and differential coverage across metagenomes. From these few metabins, a total of nine nucleomorph MAGs were characterised and manually curated (Table S2).

### A planktonic eukaryotic genomic database

The planktonic eukaryotic genomic database corresponds to 683 marine nuclear MAGs^31^ characterised from the 11 large metagenomic co-assemblies of *Tara* Oceans. We also included 705 marine plastid MAGs^9^ (also characterised from the 11 large metagenomic co-assemblies of *Tara* Oceans) and 163 reference plastid genomes from culture species (Table S5).

### Cryptophyceae genomic database

As additional context for our analyses of *Teleaulax* genomes, we created a Cryptophyceae database that covers the nuclear, nucleomorph and plastid genomes from cultured Cryptophyceae species available from NCBI at the time of our investigations (Table S5).

### Gene prediction in nucleomorph genomes from MAGs and culture

Gene prediction was performed for the nine *Teleaulax* nucleomorph MAGs, the nucleomorph genome of *Cryptomonas gyropyrenoidosa* (strain SAG 25.80)^22^ and 11 Cryptophyceae nuclear genomes using MetaEuk v5^59^ with the easy-predict workflow (“metaeuk easy-predict -s 7.5”). Protein sequences from the planktonic eukaryotic genomic database, other cellular MAGs characterised from the same 11 metagenomic co-assemblies^31,38^, the additional Cryptophyceae database and a manually curated version of MarFERReT^60^ resource (from which sequences with multiple stop codons were further removed) served as reference for the predictions.

### A concatenated gene alignment phylogeny of Cryptophyceae nucleomorphs

We used 240 Hidden Markov model (HMM) profiles of eukaryotic protein-coding genes from the PhyloFisher^61^ database with the anvi’o program “anvi-run-hmms” (default parameters; the program uses HMMER^57^ v3.1b2) to identify the corresponding genes 8 red algae reference genomes from culture (downloaded from NCBI), the nucleomorph MAGs (n=9) and nucleomorph reference genomes from the Cryptophyceae database (n=9). After considering the number of hits per HMM across the genomes, we created a collection of 33 single-copy core genes that are predominantly present as a single copy in nucleomorphs and red algal nuclear genomes alike (Table S2). We performed an alignment of each single-copy core gene across the considered genomes using MAFFT^62^ with default parameters and concatenated the alignments. Alignment sites containing more than 30% gaps were trimmed using trimAl^63^ v1.4.1 (“-gapthreshold 0.3”). Maximum likelihood trees were inferred with IQ-TREE^64^ v1.6.12 (“-m MFP -safe -alrt 1000 -bb 1000” parameters), using the ModelFinder^65^ Plus option to determine the best-fitting substitution model according to the Bayesian Information Criterion. Branch supports were calculated from 1,000 replicates each for the *Shimodaira–Hasegawa*-like approximate likelihood ratio test (SH-like aLRT) and ultrafast bootstrap (UFBoot). Nodes were considered well supported if they met SH-like aLRT ≥ 80% and UFBoot ≥ 95%, moderately supported if only one of the two thresholds was met, and poorly supported otherwise. Phylogenetic trees were visualized and rooted using anvi’o.

### Distribution of Cryptophyceae nucleomorph genomes in the sunlit oceans

We mapped 1,178 *Tara* Oceans metagenomes to our planktonic eukaryotic genomic database and to the Cryptophyceae database and determined genome-scale mean coverages (vertical coverage) and detection scores (horizontal coverage). Mapping was performed with BWA^66^ v0.7.15 (minimum identity of 95%) using a FASTA file containing 1,390 MAGs and 177 reference genomes to recruit short reads from all metagenomes. A genome was considered detected in a given filter when ≥25% of its length was covered by reads, to minimize non-specific read recruitment^67^. The number of recruited reads below this cutoff was set to 0 before calculating vertical coverage and the percentage of recruited reads (Table S3).

### Correlation between Teleaulax nucleomorph MAGs and other genomes

For each of the nine *Teleaulax* nucleomorph MAGs, a Pearson correlation coefficient of coverage across 1,178 *Tara* Oceans metagenomes was calculated against each genome of the planktonic eukaryotic genomic database. Pearson correlation coefficients were subsequently used, in the context of taxonomic assignments and phylogenies, to perform an environmental linkage connecting the nuclear, nucleomorph and plastid genomes of *Teleaulax* lineages characterised in our study.

### Gene expression of *Teleaulax* genomes in the sunlit oceans

We mapped 1,165 Tara Oceans metatranscriptomes using BWA v0.7.15 (minimum identity of 95%) against a FASTA file containing genes from the nucleomorph MAGs as well as the planktonic eukaryotic genomic database and Cryptophyceae genomic database to recruit metatranscriptomic short reads. Metatranscriptomic read recruitment was used to calculate the mean coverage (vertical coverage) of individual genes as well as genome-scale detection scores (horizontal coverage). Downstream metatranscriptomic analyses were restrained to filters in which ≥ 50% of the genes from a given genome were detected (Table S4). For *Teleaulax*, *in situ* gene expression of DNApol-III subunits was subsequently investigated across the nuclear, nucleomorph and plastid genomes based on the recruited metatranscriptomic reads.

### Culture of *Teleaulax amphioxeia* strain AND-A0710 and DNA extraction

Culture of *Teleaulax amphioxeia* strain AND-A0710 was maintained in L1 seawater medium at 20°C, without agitation, under white light at ∼150 µmol photons m^-2^s^-1^ on a 12h/12h light/dark cycle. DNA was extracted from 138 million cells using the DNAbsolute kit (Eagle Biosciences, NH, USA) following manufacturer instructions. Quantification was performed using a Qubit dsDNA HS Assay kit (Thermo Fisher Scientific) and integrity was assessed in a FemtoPulse system (Agilent). DNA was stored at 4 °C until usage.

### Library preparation and sequencing of *Teleaulax amphioxeia* strain AND-A0710

DNA fragment size selection was performed using Short Read Eliminator (PacBio, CA, USA). Selected DNA was then fragmented using the Megaruptor 3 (Diagenode, Liege, Belgium) and purified with the DNeasy PowerClean Pro Cleanup (Qiagen). Long-read DNA library was prepared with the SMRTbell prep kit 3.0 following manufacturers’ instructions and sequenced on a Revio system (PacBio). Hi-C libraries were generated from 99 million cells using the Arima High Coverage Hi-C kit (following the Animal Tissues low input protocol v01) and sequenced on a NovaSeq X instrument (Illumina) with 2x150 bp read length. In total 11.78 Gb PacBio HiFi and 228.8 Gb HiC data were sequenced to generate the assembly.

### Genome Assembly of *Teleaulax amphioxeia* strain AND-A0710

The genome of *Teleaulax amphioxeia* strain AND-A0710 was assembled using the Genoscope GALOP pipeline (https://workflowhub.eu/workflows/1200). Briefly, raw PacBio HiFi reads were assembled using Hifiasm v0.24.0-r703. Resulting contigs were screened for non-eukaryotic sequences and 73 contigs were identified as contaminants, totalling 10.131 Mb (with the largest being 5.82 Mb). Retained haplotigs (286 regions totalling 8.023 Mb) were removed using purge_dups v1.2.5 with default parameters and the proposed cutoffs. The purged assembly was scaffolded using YaHS v1.2.2 and assembled scaffolds were then curated through manual inspection using PretextView v0.2.5 to remove false joins and incorporate sequences not automatically scaffolded into their respective locations within the chromosomal pseudomolecules. During manual curation, 6 haplotypic regions and 184 contaminant sequences were removed, totalling 0.271 Mb and 5.26 Mb, respectively (with the largest being 0.091 Mb and 0.126 Mb). From these 184 removed sequences, 183 correspond to another eukaryotic species (*Thraustochytrida order*). The nuclear genome is composed of 101 chromosomes, while chromosomes 102, 103, and 104 correspond to the nucleomorph genome. The mitochondrial and plastid genomes were assembled using OATK^68^ and included in the released assembly (Table S6).

### Gene prediction in *Teleaulax amphioxeia* strain AND-A0710

Protein-coding genes in the nuclear and nucleomorph genomes of *Teleaulax amphioxeia* strain AND-A0710 were predicted using Metaeuk v5^59^ (as described in the section “Gene prediction in nucleomorph genomes from MAGs and culture”). Gene prediction for mitochondrial and plastid genomes was performed using Prodigal v2.6.3.1^69^ (default parameters).

### Average nucleotide identity between genomes

Average nucleotide identity (ANI) values between genomes were computed using skani^70^ v0.2.1 (“skani triangle”, default settings) across four comparison sets: (1) among *Teleaulax* nucleomorph MAGs; (2) between *Teleaulax* nucleomorph MAGs and Cryptophyceae reference nucleomorph genomes from culture; (3) among *Teleaulax* nuclear MAGs, (4) and among *Teleaulax* plastid MAGs. Finally, ANI was also calculated between plastid genomes of the cultured *Teleaulax amphioxeia* strains HACCP-CR01 and AND-A0710 (Table S2).

### Discovery of DNApol-III subunits in Teleaulax genomes

We predicted Pfam domains for Cryptophyceae nucleomorph genomes using the anvi’o genomic workflow, and Pfam occurrence and functional enrichment were computed across genomes using the anvi’o program “anvi-compute-functional-enrichment-across-genomes”^71^ (Table S2). This analysis identified the DNApol-III alpha subunit in nucleomorph genomes of *Teleaulax* and close relatives within the Pyrenomonadales. We subsequently performed a large-scale functional annotation of proteins using InterProScan^72^ v5.69-101.0 with the options “-dp -dra -goterms - iprlookup”, on the *Teleaulax* MAGs (nuclear, nucleomorph, plastid) and four genomes of *Teleaulax amphioxeia* strain AND-A0710. This analysis identified additional DNApol-III subunits in the genomic landscape of *Teleaulax*.

### A global search for eukaryotic DNApol-III on NCBI

We BLASTP-searched protein sequences corresponding to the four DNApol-III subunits from 19 cyanobacterial orders and identified in *Teleaulax* genomes (nucleomorph, nuclear and plastid) against the NCBI ClusteredNR database (downloaded in November 2025 https://ftp.ncbi.nlm.nih.gov/blast/db/experimental/), restricting searches to Eukaryota taxid 2759 (default settings and maximum target sequences set to 5,000 hits). For each query, non-redundant hits with the highest bitscore per query were retained (minimum e-value < 1×10^-5^) (Table S9). The hits corresponded to genomes of green algae and red algae. We downloaded the predicted proteins from the corresponding genomes when available in NCBI, resulting in a database of 22 nuclear genomes and 15 corresponding plastid genomes (Table S5).

### Cyanobacterial genomic database

The Cyanobacteria genomic database corresponds to a previously published global database of cyanobacterial genomes^73^, from which we excluded genomes shorter than 1 Mb. Gene prediction was performed using Prodigal v2.6.3.1^69^ (default parameters). We identified DNApol-III alpha subunit using hmmsearch^57^ (HMMER v3.4; “--cut_ga”) and the reference Pfam HMM profiles PF17657. We used a e-value ≤ 1×10^-40^ and only retained proteins ranging from 800 to 2,500 amino acids. WE also excluded proteins containing multiple ambiguous residues (“X”). We used CD-HIT^74^ v4.8.1 (“-c 0.80”) to select one representative alpha subunit sequence per protein cluster at the 80% identity level. Cyanobacterial genomes corresponding to these representative sequences were selected and genomes with completion below 50% or redundancy above 25% were excluded, resulting in a final database of 533 genomes spanning all major cyanobacterial lineages (Table S10).

### Identification of DNApol-III subunits in eukaryotic and cyanobacterial genomes

A global hmmsearch^57^ (HMMER v3.4; “--cut_ga”) for DNApol-III subunits was performed, at the protein level, for the Cryptophyceae genomic database, planktonic eukaryotic genomic database, cyanobacterial genomic database, other eukaryotic genomes (nuclear and plastid) with DNApol-III hits from NCBI (see “A global search for eukaryotic DNApol-III on” section), and finally Archaeplastida and Chlorarachniophyte reference nuclear genomes (107 Chlorophyta; 46 Rhodophyta; 17 Streptophyta; 1 Glaucophyta; gene calling with MetaEuk v5^59^ when reference proteins were not available) and plastid genomes (215 Chlorophyta, 234 Rhodophyta, 30 Streptophyta and 4 Glaucophyta; gene calling with Prodigal v2.6.3.1^69^ when reference proteins were not available) available from NCBI’s Genome resource (https://www.ncbi.nlm.nih.gov/datasets/genome/) (Table S5). For the alpha and beta subunits, we used the reference Pfam HMM profiles PF17657 (alpha) and PF00712 (beta). For alpha, we used a e-value ≤ 1×10^-40^ and only retained proteins ranging from 800 to 2,500 amino acids. For beta, we used a e-value ≤ 1×10^-20^ and only retained proteins ranging from 280 to 750 amino acids. For the tau/gamma subunit, we built a custom HMM profile. We first extracted all the bacterial proteins annotated with InterPro entry IPR012763 (n=20,960; accessed in November 2025), retained those ranging from 300 to 800 amino acids, and created a non-redundant database of 13,056 proteins (95% identity) using CD-HIT v.4.8.1^74^. The resulting sequences were aligned using MAFFT^62^ with default parameters, and the alignment was used to build the HMM profile of tau/gamma with hmmbuild^57^ (HMMER v3.4). During our global search for tau/gamma in the eukaryotic and cyanobacterial genomes, we used a e-value ≤ 1×10^-25^ and only retained proteins ranging from 350 to 1,500 amino acids. Finally, for the epsilon subunit, we also built a custom HMM profile. We first extracted cyanobacterial protein sequences retrieved from NCBI (n=66,961; accessed in November 2025) using the query “DNA polymerase III epsilon subunit”, retained those ranging from 200 to 350 amino acids, and created a non-redundant database of 12,134 proteins (95% identity) using CD-HIT v.4.8.1^74^. The resulting sequences were aligned using MAFFT^62^ with default parameters, and the alignment was used to build the HMM profile of epislon with hmmbuild^57^ (HMMER v3.4). During our global search for epsilon in the eukaryotic and cyanobacterial genomes, we used a e-value ≤ 1×10^-10^ and only retained proteins ranging from 150 to 500 amino acids. For all four subunits, in rare cases where multiple proteins corresponding to the same subunit were detected in a genome, the longest sequence was retained for downstream analyses. In the case of DNApol-III alpha subunit identified in the reference genome of *Rhinomonas nottbeckii* (GCA_965120255.1), we determined using protein-level taxonomic assignments that the corresponding contig corresponded to its nucleomorph genome.

### Phylogenetic analysis of DNApol-III subunits

Single protein phylogenies were computed for the alpha, beta and tau/gamma DNApol-III subunits using sequences from Cryptophyceae, cyanobacteria and additional eukaryotic homologs. For these subunits, Actinomycetota and Chloroflexia sequences were included for rooting (as previously described^75^) (Table S10). For the epsilon subunit, in addition to homologs identified in eukaryotes and cyanobacteria, additional protein homologs identified from 1,778 marine bacterial MAGs^38^ using HMM searches with the epsilon profile described above were included, and the tree was rooted using PolC proteins, as previously described^75^. For all four subunits, protein sequences were aligned using MAFFT v7.526^62^ with the L-INS-i algorithm and default parameters. Trimming and tree inference followed the same protocol as described in the section “A concatenated gene alignment phylogeny of Cryptophyceae nucleomorphs”. Trees were visualised and rooted using anvi’o, and long branches were iteratively removed manually.

### Phylogenetic analysis of DNA polymerase family B in Archaeplastida

DNA polymerase family B (DNApolB) protein homologs were identified from nuclear genomes of Archaeplastida with hmmsearch^57^ (HMMER v3.4; “--cut_ga”) using the Pfam HMM profile PF00136. Protein sequences were aligned using MAFFT v7.526^62^ with the L-INS-i algorithm and default parameters. Alignment trimming and phylogenetic inference followed the same protocol as described above (see section “A concatenated gene alignment phylogeny of Cryptophyceae nucleomorphs”). Phylogenetic tree was visualised and rooted using anvi’o.

### Identification and comparative expression analysis of transcripts encoding DNApol-III subunits in *Teleaulax amphioxeia* and *Mesodinium rubrum*

Transcript abundance estimates for *Teleaulax amphioxeia* SCCAP K-1837 and *Mesodinium rubrum* MBL-DK2009 across 27 samples were obtained from a previous study^51^. These transcripts were reassigned by BLASTN (e-value ≤ 10^-5^) to the nuclear, nucleomorph, plastid and mitochondrial genomes of *T. amphioxeia* strain AND-A0710 to infer the genomic type of each transcript and identify transcripts associated with DNApol-III subunits. *M. rubrum* transcripts were independently translated in all six reading frames and annotated with InterProScan^72^ v5.69-101.0 to identify transcripts encoding the DNApol-III subunits. Transcript abundances were then normalised separately for the *T. amphioxeia* and *M. rubrum* transcript sets by scaling each sample to 1 million reads, and normalised values were used for downstream expression analyses.

### Prediction of signal peptides and subcellular localization of DNApol-III subunits

Genes of DNApol-III subunits found in eukaryotes were analyzed using SignalP v6.0^76^ (organism types set to “Eukarya” and “Other”, respectively; model mode Slow), TargetP v2.0^77^ (organism types “Plant” and “Non-plant”, respectively) and DeepLoc v2.1^78^ (default settings). For DNApol-III genes lacking detectable plastid-targeting signals, upstream genomic segments (900 nucleotides on the N-terminal side of the predicted protein) were added to overcome possible errors with the gene prediction models. These segments were retained only when they did not overlap neighbouring predicted genes on the same frame, and their reading frame was determined by BLASTP searches against the NCBI ClusteredNR database. With this approach, signal peptides could be found in multiple upstream genomic segments, always in the 300 nucleotides closets to the start of the predicted gene.

## Supplementary Tables

**Table S1: Description of 1,178 Tara Oceans metagenomes and 1,165 metatranscriptomes.** This table reports main metadata for Tara Oceans metagenomes and metatranscriptomes, including sampling depth, oceanic region, filter size fractions and sequencing type.

**Table S2: Genomic statistics and functional annotation of Cryptophyceae nucleomorph genomes.** This table reports genomic statistics for Cryptophyceae nucleomorph genomes and average nucleotide identity (ANI) values between environmental linked *Teleaulax* genomes and the cultured *Teleaulax amphioxeia* strain AND-A0710. It also lists the 33 conserved single-copy core genes selected based on their occurrence across *Teleaulax* nucleomorph MAGs, nucleomorph genomes from cultured Cryptophyceae and eight cultured red algal genomes. Finally, it summarises Pfam annotation occurrence and functional enrichment statistics comparing *Teleaulax* nucleomorph genomes with previously sequenced Cryptophyceae nucleomorph genomes.

**Table S3: Biogeography and environmental linkage.** This table reports the biogeographic signal of genomes from the nucleomorph MAG database, the planktonic eukaryotic genomic database and the additional Cryptophyceae database across 1,178 Tara Oceans metagenomes, including coverage statistics by depth, size fraction and oceanic region. It also provides Pearson correlation coefficients of coverage across Tara Oceans metagenomes calculated between nucleomorph MAGs and genomes from the eukaryotic genomic database. This table reports inferred genomic ratios of environmental linkage between nucleomorph, plastid and nuclear genomes.

**Table S4: In situ gene expression.** This table reports mapped metatranscriptomic reads for nucleomorph genomes (MAGs and Cryptophyceae reference genomes from culture) and in situ expression levels of the four DNApol-III subunit genes (alpha, epsilon, beta and tau/gamma) across environmental linked *Teleaulax* genomes from 1,140 Tara Oceans metatranscriptomes.

**Table S5: Eukaryotic genomic resource.** This table compiles all eukaryotic genomes analysed in this study with associated metadata (accession, genomic compartment, taxonomy and genomic statistics). The occurrence of the four DNApol-III subunits (alpha, epsilon, beta and tau/gamma) is indicated for each genome.

**Table S6: Genomic statistics of *Teleaulax amphioxeia strain AND-A0710*.** This table reports chromosome level genomic statistics across the different genomes of *T. amphioxeia* strain AND-A0710 and reports the presence or absence of DNApol-III subunits in each genome.

**Table S7: Signal peptide and subcellular localization predictions of DNApol-III subunits.** This table reports the identification of signal peptide and subcellular localization predictions in genes corresponding to DNApol-III subunits and found in eukaryotic genomes.

**Table S8:** Identification and expression analyses of DNApol-III subunit associated transcripts in *Teleaulax amphioxeia* and *Mesodinium rubrum*. This table describes our BLASTN-based transcript reassignment to *Teleaulax amphioxeia* strain AND-A0710 genomes, InterProScan based annotation of *Mesodinium rubrum* transcripts, normalized expression values, and statistical analyses of DNApol-III subunit-associated transcripts.

**Table S9: A global search for eukaryotic DNApol-III subunits.** This table reports BLASTP hits against the NCBI ClusteredNR database and retains non-redundant eukaryotic hits with the highest bitscore per subunit query.

**Table S10: Bacterial genomic database.** This table compiles genomic databases from Cyanobacteria, Actinomycetota and Chloroflexota, with accession information, taxonomic assignment and genomic statistics.

**Table S11: Functional annotations of Cryptophyceae genomes.** This table reports InterProScan functional annotation for genes from environmental linked *Teleaulax* genomes, the cultured *Teleaulax amphioxeia* strain AND-A0710, and additional reference Cryptophyceae genomes. For each gene, associated InterProScan accessions and functional descriptions are reported, and genes encoding DNApol-III subunits are indicated.

## Supplementary Figures

**Figure S1:**
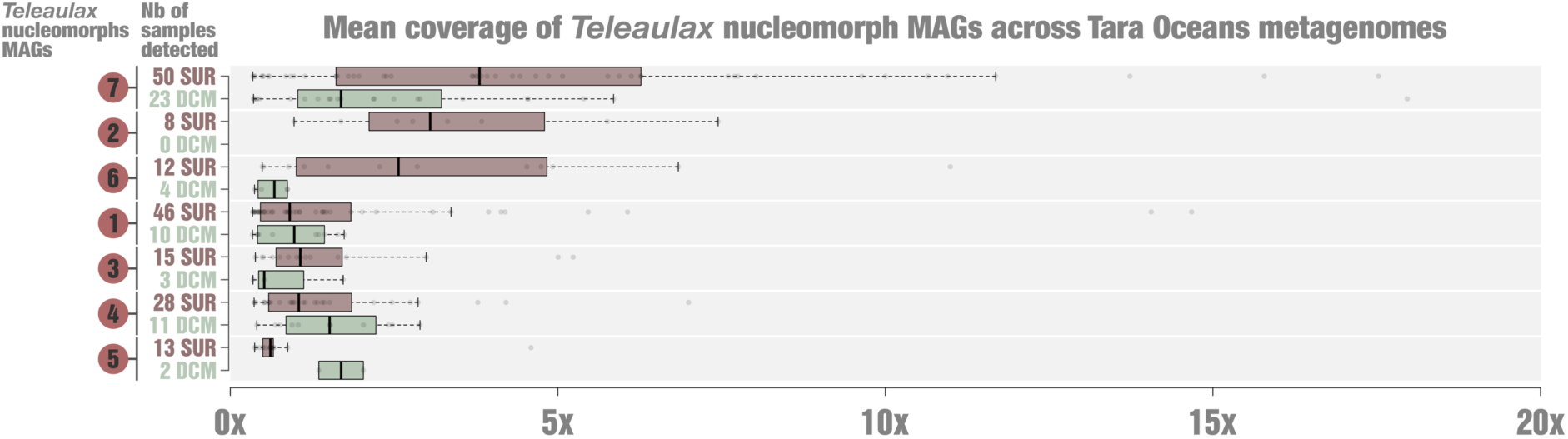
Environmental signal of *Teleaulax* nucleomorph MAGs in the sunlit oceans. For each *Teleaulax* nucleomorph MAG, boxplots represent mean coverage across *Tara* Oceans metagenomic samples collected at the surface (SUR) and deep chlorophyll maximum (DCM). Only samples in which the genome was detected (coverage > 0) are shown. Center lines in boxplots indicate the medians, box limits correspond to the 25^th^ and 75^th^ percentiles, whiskers extend 1.5 times the interquartile range and outliers are plotted individually. The number of detected SUR and DCM samples is reported for each MAG.

**Figure S2:**
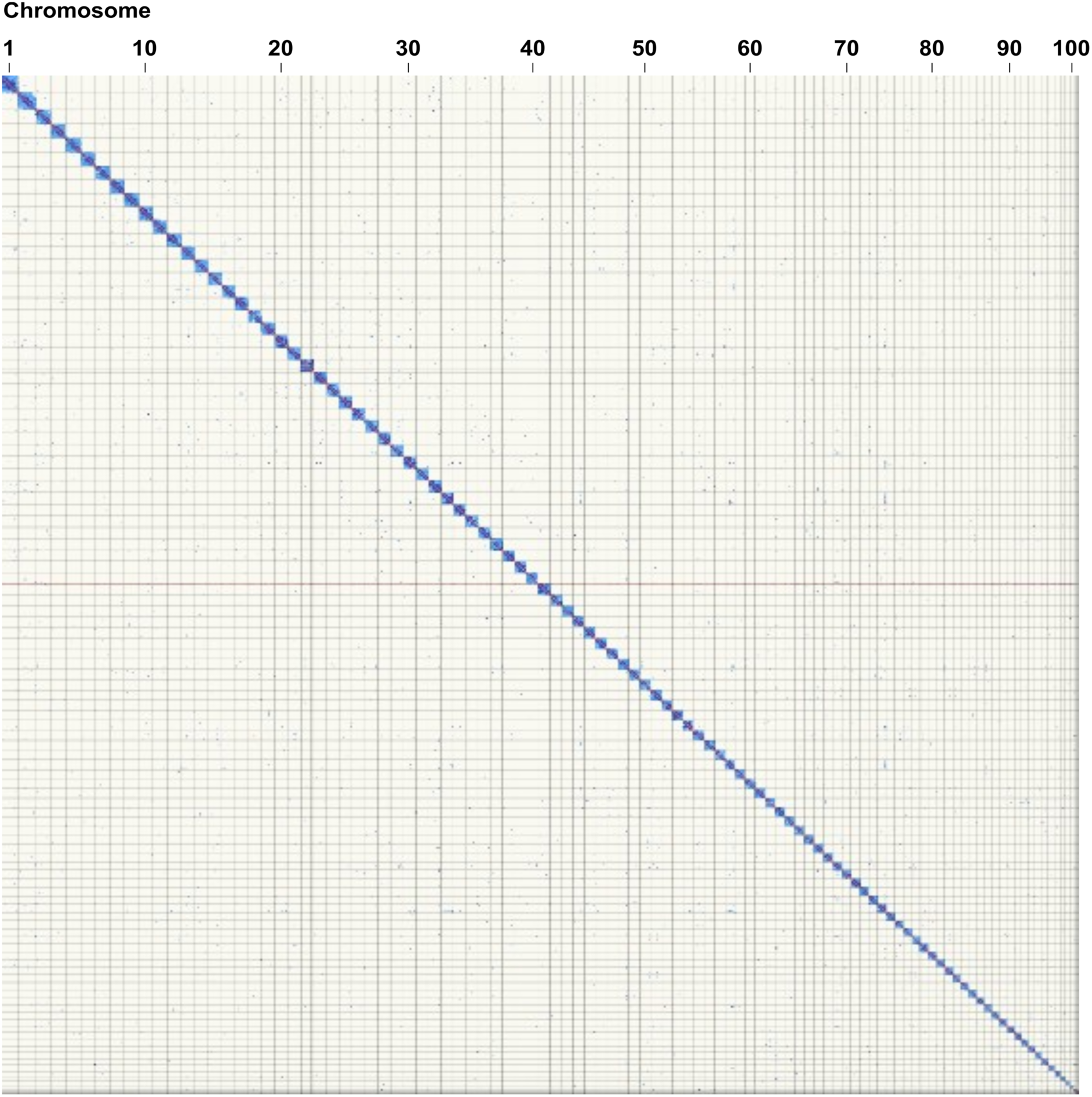
Hi-C contact map of the *Teleaulax amphioxeia* strain AND-A0710 genome assembly. Hi-C contact map of interaction frequencies among assembled genomic regions of *Teleaulax amphioxeia* strain AND-A0710 assigned to the nuclear, nucleomorph, plastid and mitochondrial genomes. Blue regions indicate higher-frequency Hi-C interactions between pairs of genomic regions.

**Figure S3:**
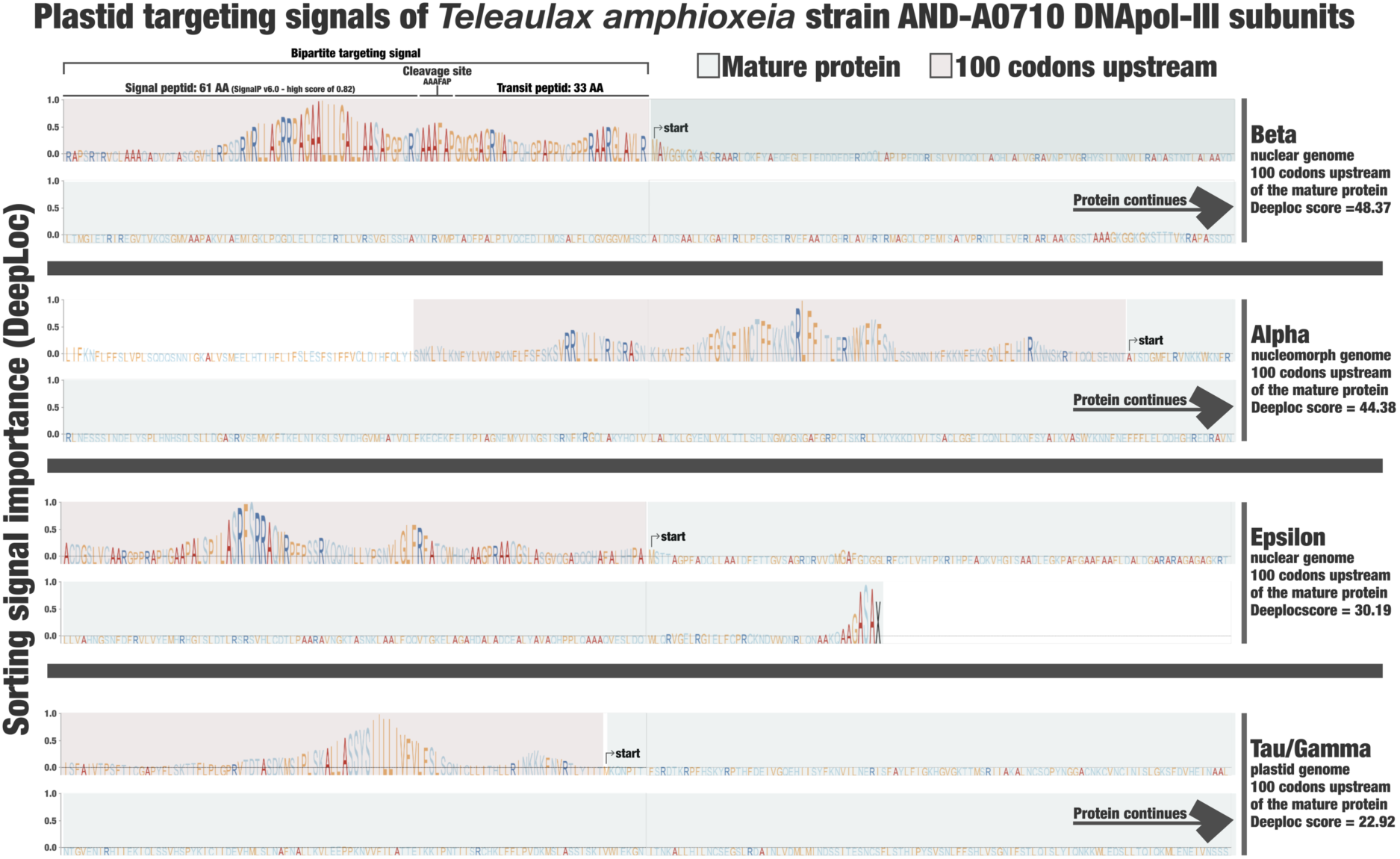
Plastid targeting signals of *Teleaulax amphioxeia* strain AND-A0710 DNApol-III subunits. The figure describes amino-acid level plastid targeting signal (“sorting signal importance” in the y-axis) as predicted by DeepLoc, for the DNApol-III beta, alpha, epsilon and tau/gamma subunits of *Teleaulax amphioxeia* strain AND-A0710. Red areas correspond to the upstream genomic segments (300 nucleotides on the N-terminal side) of predicted protein. Green areas correspond the beginning of the mature protein. For the beta subunit, we identified a bipartite plastid-targeting signal including a 61 amino acid signal peptide predicted using SignalP.

**Figure S4:**
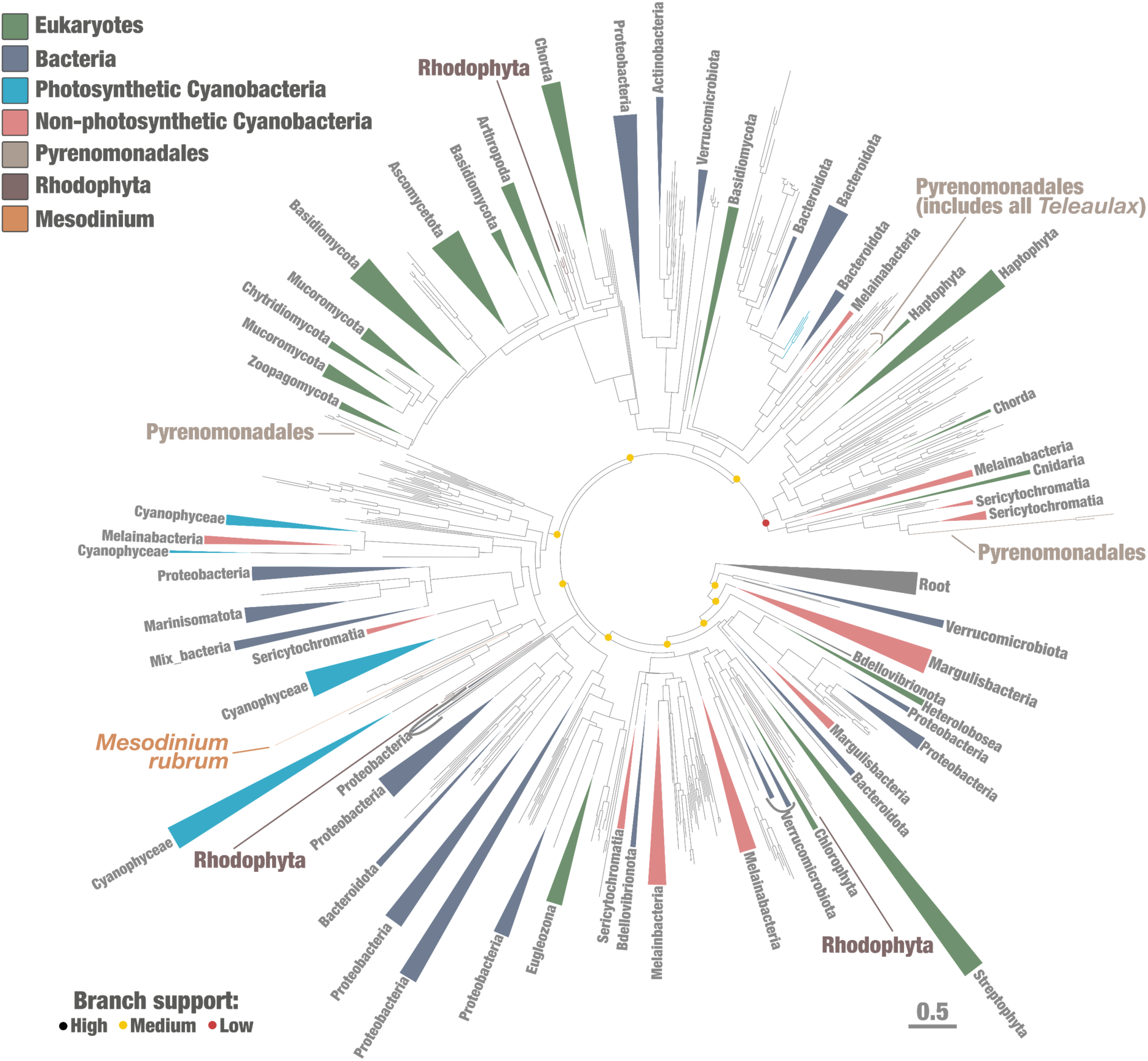
Single-protein phylogeny of the DNA polymerase III epsilon subunit. Maximum-likelihood phylogenetic tree of the DNApol-III epsilon subunit (251 amino acid positions; Q.PFAM+R10) including 3,138 sequences (2,048 Bacteria and 1090 Eukarya). The tree was rooted using PolC proteins as previously described^75^. Highlighted phylogenetic supports (dots) were considered high (SH-like aLRT ≥80 and UFBoot≥95, in black), medium (SH-like aLRT ≥80 or UFBoot≥95, in yellow) or low (SH-like aLRT<80 and UFBoot<95, in red). Trees were decorated with additional information and visualised with anvi’o. Scale bar indicates the number of substitutions per site for each tree.

**Figure S5:**
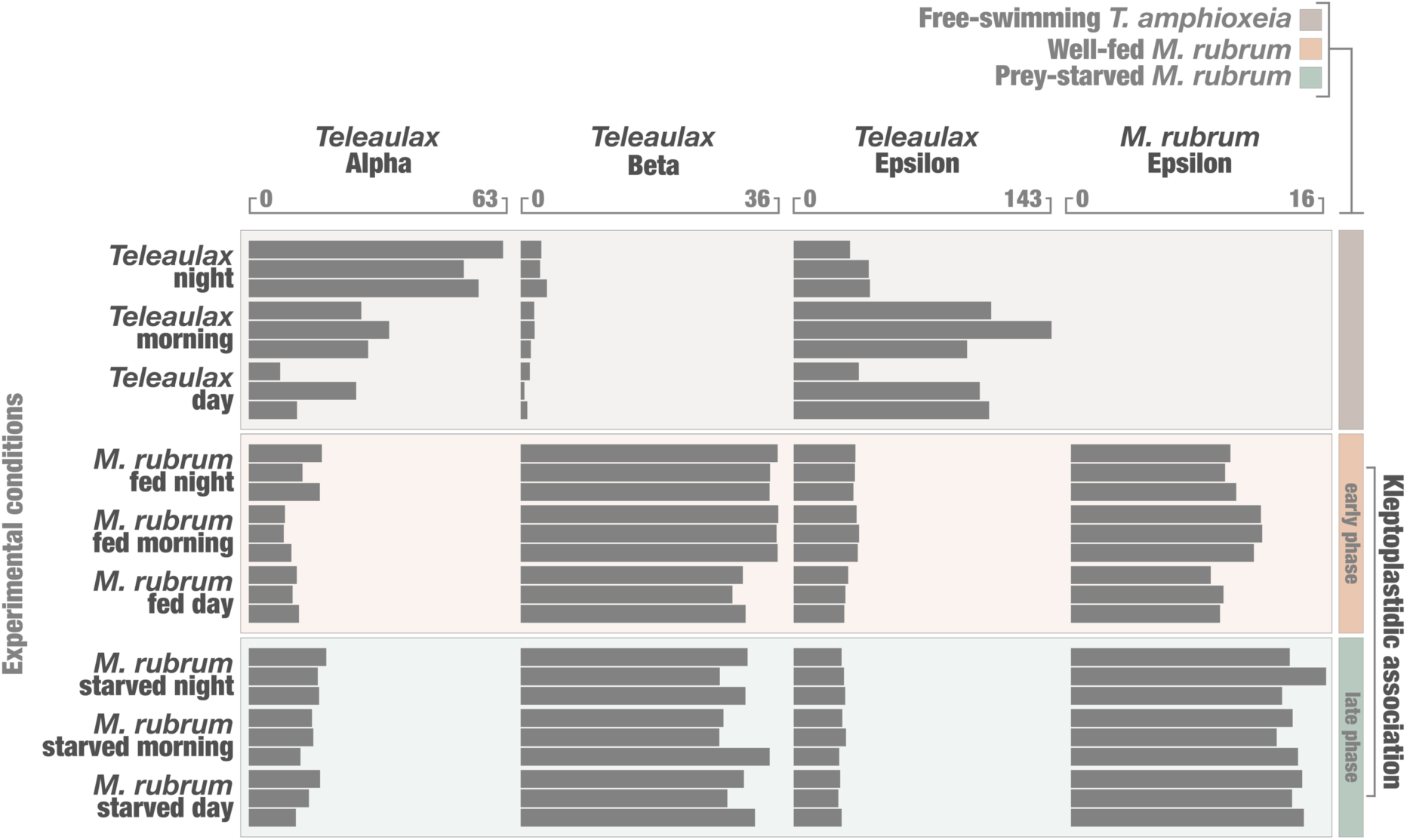
Expression of DNApol-III subunit associated transcripts. Expression levels of transcripts associated with DNApol-III subunits in *T. amphioxeia* (alpha, beta and epsilon) and *M. rubrum* (epsilon) across nine experimental conditions (free-swimming *T. amphioxeia*, well-fed *M.* rubrum and prey-starved *M. rubrum* at night, morning and day). Expression values were normalized to 1 million reads. Bars represent individual transcript abundances for each subunit and condition.

**Figure S6:**
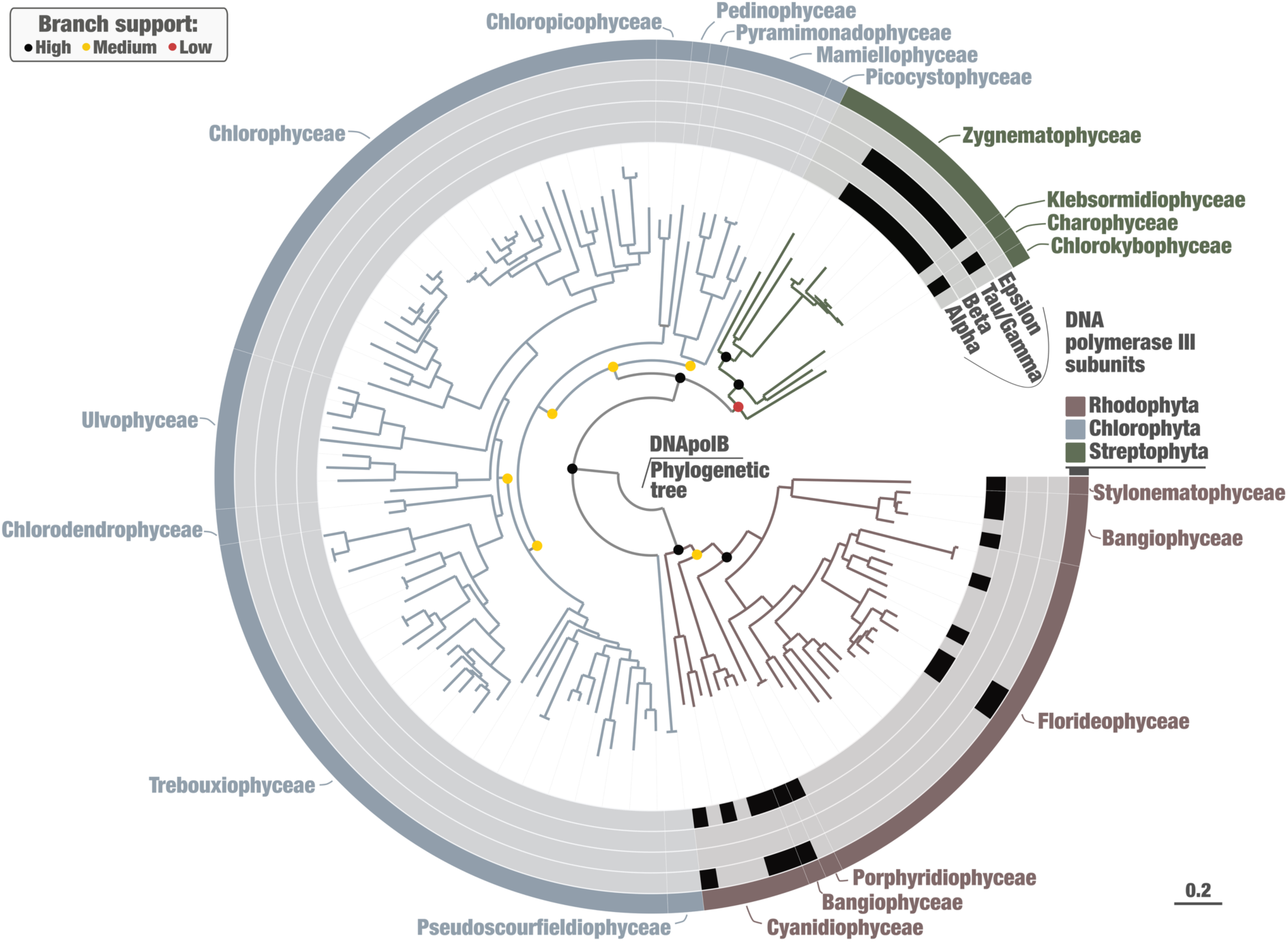
Single-protein DNA polymerase B phylogeny and DNApol-III occurrence in Archaeplastida (Rhodophyta, Chlorophyta and Streptophyta). Maximum-likelihood phylogenetic tree of DNA polymerase B (1,091 amino acid positions, Q.YEAST+F+I+R7 model) including 139 sequences (35 Rhodophyta, 90 Chlorophyta and 14 Streptophyta). Highlighted phylogenetic supports (dots in the trees) were considered high (SH-like aLRT ≥80 and UFBoot≥95, in black), medium (SH-like aLRT ≥80 or UFBoot≥95, in yellow) or low (SH-like aLRT<80 and UFBoot<95, in red). The scale bar represents 0.2 substitutions per site. The tree was decorated with Archaeplastida class-level and visualized using anvi’o.

**Figure S7:**
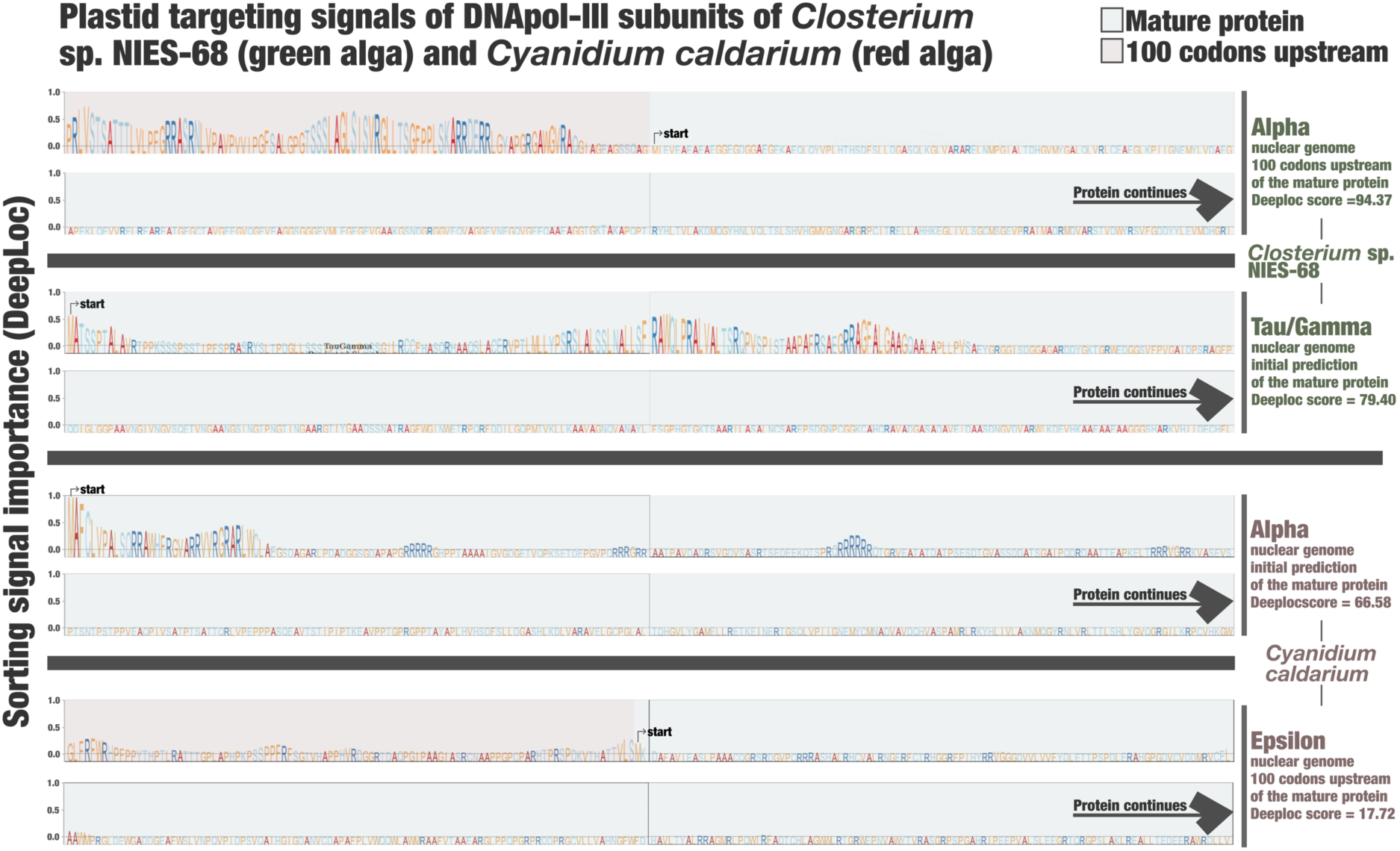
Plastid targeting signals of DNApol-III subunits of *Closterium* sp. NIES-68 (green alga) and *Cyanidium caldarium* (red alga). The figure describes amino-acid level plastid targeting signal (“sorting signal importance” in the y-axis) as predicted by DeepLoc, for the DNApol-III alpha and tau/gamma subunits of *Closterium* sp. NIES-68 and the alpha and epsilon subunits of *Cyanidium caldarium*. Red areas correspond to the upstream genomic segments (300 nucleotides on the N-terminal side) of predicted protein. Green areas correspond the beginning of the mature protein.

**Figure S8:**
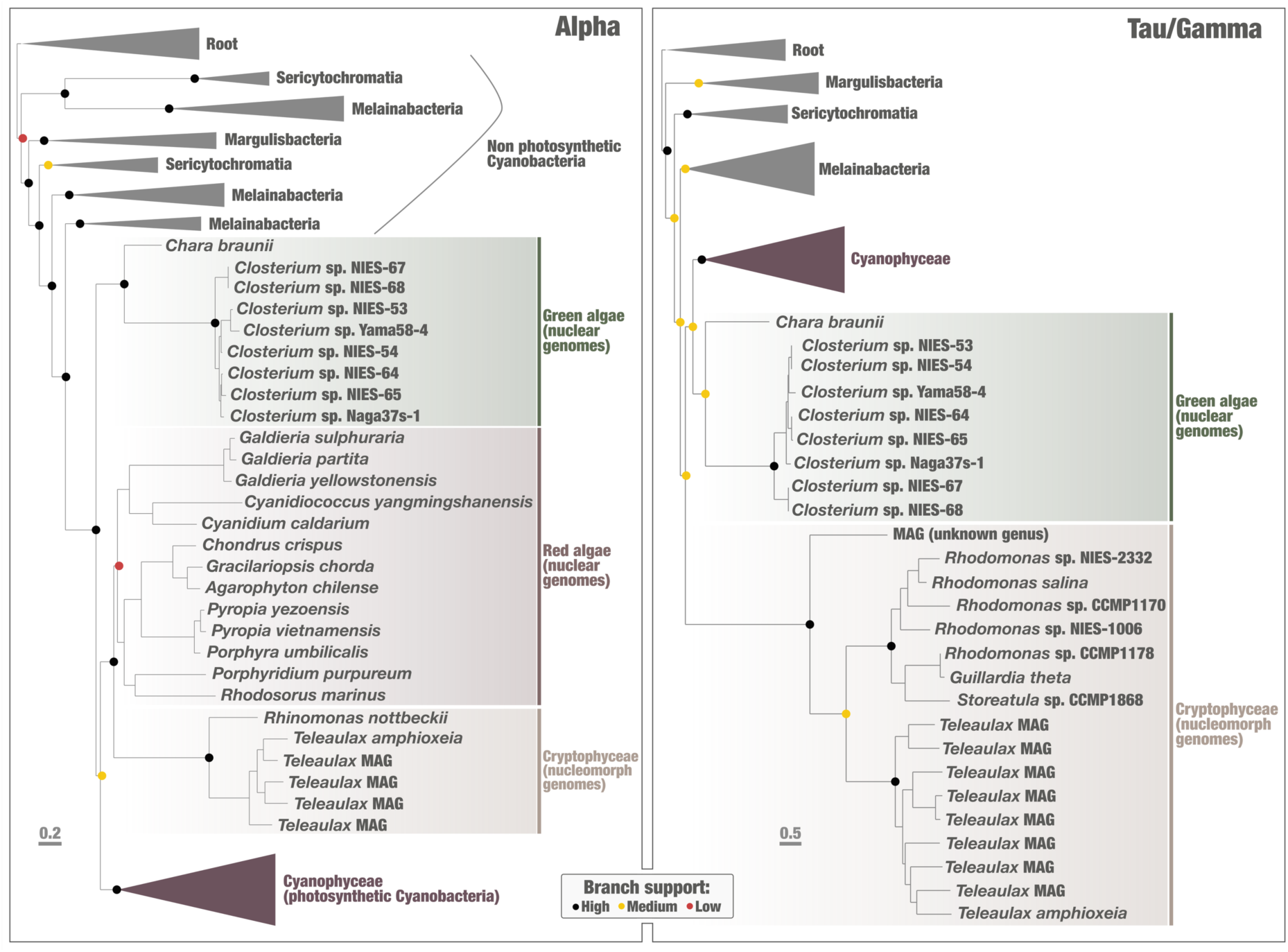
Single-protein phylogenies of the DNApol-III alpha and tau/gamma subunits with Archaeplastida including *Chara braunii*. Left panel displays a maximum-likelihood phylogenetic tree of the DNApol-III alpha subunit (1,178 amino-acid positions, LG+I+R10 model), including 568 sequences (495 Cyanobacteria, 28 Eukarya, 17 Chloroflexia and 28 Actinomycetota). Right panel displays a maximum-likelihood phylogenetic tree of the DNApol-III tau/gamma subunit (697 amino-acid positions, Q.PFAM+F+I+R10 model), including 451 sequences (397 Cyanobacteria, 26 Eukarya and 28 Actinomycetota). Highlighted phylogenetic supports (dots in the trees) were considered high (SH-like aLRT ≥ 80 and UFBoot ≥ 95, in black), medium (SH-like aLRT ≥ 80 or UFBoot ≥ 95, in yellow) or low (SH-like aLRT < 80 and UFBoot < 95, in red). The alpha tree was rooted with Chloroflexia and Actinomycetota, while the tau/gamma tree was rooted with Actinomycetota. Scale bars indicate the number of substitutions per site for each tree. Trees were decorated with additional layers and visualised with anvi’o.

## Data availability

All data generated in this study are publicly available via Zenodo at https://zenodo.org/records/20328056. These include (1) contigs and predicted proteins for all the Cryptophyceae genomes (nuclear, nucleomorph, plastid) used in our analyses (including the genome of *Teleaulax amphioxeia* strain AND-A0710 *with* BioSample SAMEA120642984 and assigned to Tree of Life ID (ToLID) ‘uyTelAmph1’), (2) contigs and predicted proteins for all the Archaeplastida genomes containing the DNApol-III alpha and tau/gamma subunits (nuclear, plastid), (3) and contigs and predicted proteins for a genomic database of Cyanobacteria^73^ that we curated in this study. In addition, the associated sequencing data for *Teleaulax amphioxeia* strain AND-A0710 is available from ENA under project PRJEB104650. Our data availability also includes (4) the concatenated alignment of 33 single-copy core genes used for Cryptophyceae nucleomorph phylogeny with the corresponding maximum-likelihood tree, (5) the alpha, beta, tau/gamma and epsilon proteins used for the global search for eukaryotic DNApol-III on NCBI, (6) HMM profiles (four DNApol-III subunits and DNApolB), (7) all predicted proteins for the DNApol-III subunits we found in eukaryotic genomes, (8) metadata, protein alignment and trees corresponding to our phylogenetic analyses of DNApol-III subunits, (9) the six-frame translated protein dataset derived from the *Mesodinium rubrum* gene assembly^51^ and (10) metadata, protein alignment and phylogenetic tree corresponding to our Archaeplastida DNApolB phylogenetic analysis. Finally, the data also includes all supplementary tables.

## Acknowledgements

Our survey was enabled by the substantial metagenomic sampling and sequencing efforts that have been generated and made publicly available in recent years. Tara Oceans (which includes the Tara Oceans and Tara Oceans Polar Circle expeditions) would not have been possible without the leadership of the Tara Oceans Foundation and the sustained support of 23 institutes (https://oceans.taraexpeditions.org/). We also acknowledge the commitment of the CNRS and Genoscope/CEA. We thank Rossana Sussarellu, Thomas Lacour, Gregory Carrier from GENALG lab at Ifremer, Nantes for providing the Teleaulax amphioxeia strain AND-A0710 culture and we are grateful to Adrien Thurotte for establishing the culture and to Karine Labadie, Corinne Cruaud, Jean-Marc Aury, Caroline Menguy and Benjamin Istace, for sequencing and assembly. We also thank Ulysse Guyet for contribution to the complete genomic resource and Morgan Gaïa for phylogenetic analyses. We are also grateful to Igor S. Pessi for providing access to a global database of cyanobacterial genomes. Some computations were performed using TGCC computing facility in France with funding from France Génomique. T. A. was funded by the Doctoral School “Structure and Dynamics of Living Systems” of Paris-Saclay University. JMA was supported by an NSERC Discovery Grant. FB’s research is supported by grants from the European Research Council (ERC consolidator grant 101044505), the Swedish Research Council VR (2025-04150), and the Olle Engkvists Stiftelse. Upon publication, this manuscript will have a contribution number from the collection of *Tara* Oceans publications.

